# Cross-dataset evaluation of deep learning models for plant pest and disease diagnosis

**DOI:** 10.1101/2024.10.07.617111

**Authors:** Jianqiang Sun

## Abstract

Deep learning models have shown significant potential for plant pest and disease (PPD) diagnosis; however, their real-world effectiveness is often limited by variability between datasets, where models trained on one dataset perform poorly on others collected under different conditions. In this study, I evaluated the cross-dataset generalization of widely used deep learning architectures, including ResNet, EfficientNet, Inception, and MobileNet, across multiple tomato pest and disease datasets. As expected, models trained and tested on the same dataset achieved high performance. However, substantial performance degradation occurred when these models were tested on different datasets, highlighting the challenges posed by dataset variability. This trend was consistent across all evaluated architectures, indicating that changing the model architecture alone is insufficient to address these issues. The findings emphasize the need for more diverse and representative datasets to better capture variability in agricultural data and enhance the practical deployment of deep learning models for PPD diagnosis.

## Introduction

Machine learning, particularly deep learning, has demonstrated significant potential in addressing complex real-world problems, including agricultural applications. However, a major barrier to its widespread adoption is the challenge of generalization, wherein models trained under specific conditions often perform poorly when applied to novel, unobserved scenarios [1–3]. This raises important concerns about the robustness of machine learning model, as real-world conditions frequently differ from controlled experimental settings.

Plant pest and disease (PPD) represent a major threat to global agriculture, and deep learning-based approaches have been extensively explored as potential diagnostic tools. Although numerous studies have investigated deep learning for PPD diagnosis [4–6], most rely on datasets collected under controlled laboratory conditions [7–9] or from constrained field environments that cover a limited range of PPD types [10–12]. For example, the widely used PlantVillage (PV) dataset [13] consists of images captured against uniform backgrounds. In contrast, real agricultural environments present diverse challenges such as variable lighting conditions, inconsistent plant orientations, and heterogeneous backgrounds. Consequently, models trained on such curated datasets often display artificially high performance that is believed or suspected to generalize poorly to practical applications.

To investigate the effects of data variability, this study evaluates the degradation in recognition performance when deep learning models are trained and tested across multiple representative tomato pest and disease datasets. The aim is to provide empirical evidence of how dataset variability influences model performance and to emphasize the need for more diverse and representative datasets to improve the real-world applicability of deep learning models in agriculture.

## Methods

### Datasets

Four tomato PPD image datasets, PlantVillage (PV) [13], Tomato-Village (TV) [14], PlantDoc [15], and Tomato Leaf Disease (TLD) [16], were selected for model evaluation. The PV dataset is widely used for developing deep learning algorithms targeting PPD. It comprises images of individual leaves detached from the plant and photographed against a uniform background. Of its 38 categories encompassing various plant species and diseases, 10 relevant to tomato pests and diseases were used in this study: healthy (HL), bacterial spot (BS), early blight (EB), late blight (LB), leaf mold (LM), tomato mosaic virus (MV), spider mites (SM), *Septoria* leaf spot (SS), target spot (TS), and yellow leaf curl virus (YV).

The TV dataset primarily contains images resembling those in PV, isolated tomato leaves on a uniform background, along with a smaller subset captured under field conditions. It includes five tomato disease categories (HL, EB, LB, leaf miner (MN), and spotted wild virus (SW)) and three categories representing physiological disorders. Categories corresponding to physiological disorders were excluded from this study.

The PlantDoc and TLD datasets include images sourced from PV, online repositories, digitized presentation slides, and field settings. Due to considerable overlap between the two, they were merged and treated as a single dataset, hereafter referred to as TLD. To avoid redundancy, images originally sourced from PV were manually removed. The final category set for the merged dataset included HL, BS, EB, LB, LM, MN, MV, SM, SS, and YV.

To ensure consistency and sufficient data representation across classes, categories with fewer than 50 images were excluded from each dataset, enhancing the reliability of comparative analysis. Uniform manifold approximation and projection (UMAP) [17] was applied to visualize the distribution of image features across datasets. Specifically, 50 images from each category per dataset were randomly sampled and resized to 224 × 224 pixels. Images were first transformed using principal component analysis, with the top 128 principal components used as input to UMAP for two-dimensional reduction. UMAP parameters were set to `n_neighbors = 10` and `min_dist = 0.1`.

### Deep learning architectures

Four widely adopted deep learning architectures, ResNet [18], Inception v3 [19], MobileNet v3 [20], and EfficientNet v2 [21], were selected for evaluation based on their high citation rates, demonstrated performance across datasets, and relevance to real-world agricultural applications. All architectures were implemented using the PyTorch library [22], specifically employing the `resnet18`, `inception_v3`, `mobilenet_v3_small`, and `efficientnet_v2_s` classes from the `torchvision.models` module. Each model was initialized with default pretrained weights from the ImageNet dataset [23].

### Cross-dataset performance evaluation

Model evaluations were conducted using all combinations of the four architectures in a five-fold cross-validation framework. For example, to assess ResNet performance on the PV dataset, the model was trained on 80% of randomly sampled PV images and validated on the remaining 20%. To evaluate cross-dataset generalization, the trained model was also validated on the 20% of randomly sampled images from the other datasets (e.g., TV and TLD). Each evaluation process was repeated five times to ensure robustness, and the average performance metrics were calculated for each model–dataset combination. Confusion matrices were generated for ResNet models trained on the PV dataset and validated on PV, TV, and TLD using results from the first validation fold.

### Performance metrics

Model performance was evaluated using the macro-averages of precision, recall, and F1 score. These metrics were calculated as follows: precision = TP / (TP + FP), recall = TP / (TP + FN), and F1 score = (2 × precision × recall) / (precision + recall), where TP, FP, and FN denote the true positive, false positive, and false negative counts, respectively.

## Results and Discussion

### Dataset characteristics

The PV dataset contains substantially more images than the other datasets, whereas the TV and TLD datasets include relatively fewer images (Table 1). In terms of disease category coverage, PV and TLD encompass a broad spectrum of categories. Only three categories, HL, EB, and LB, are common across all datasets.

**Table 1.** **Number of images per category in the** PlantVillage (PV), Tomato-Village (TV), and Tomato Leaf Disease (TLD) datasets used for model evaluation. Values represent the number of images corresponding to each disease or condition in the respective datasets.

| Category | PV | TV | TLD |
| --- | --- | --- | --- |
| healthy (HL) | 1,591 | 226 | 212 |
| bacterial spot (BS) | 2,127 | - | 110 |
| early blight (EB) | 1,000 | 62 | 423 |
| late blight (LB) | 1,909 | 125 | 350 |
| leaf mold (LM) | 952 | - | 290 |
| leaf miner (MN) | - | 1,824 | 380 |
| mosaic virus (MV) | 373 | - | 116 |
| two spotted spider mites (SM) | 1,676 | - | 54 |
| Septoria leaf spot (SS) | 1,771 | - | 288 |
| spotted wild virus (SW) | - | 611 | - |
| target spot (TS) | 1,404 | - | - |
| yellow leaf curl virus (YV) | 5,357 | - | 145 |
| Total | 18,160 | 2,848 | 2,368 |
| Categories | 10 | 5 | 10 |

UMAP visualizations revealed that images clustered predominantly by dataset rather than by disease type (Figure 1), highlighting the challenge of generalization across datasets: models trained on images collected under one set of conditions often perform poorly on images acquired under different conditions. The weak separation by disease type likely reflects intra-dataset compositional uniformity. For instance, PV images typically feature a single leaf against a monochromatic background, which may dominate the feature space captured by UMAP. Consequently, subtle disease symptoms are masked by image composition, complicating accurate disease classification.

**Figure 1.**
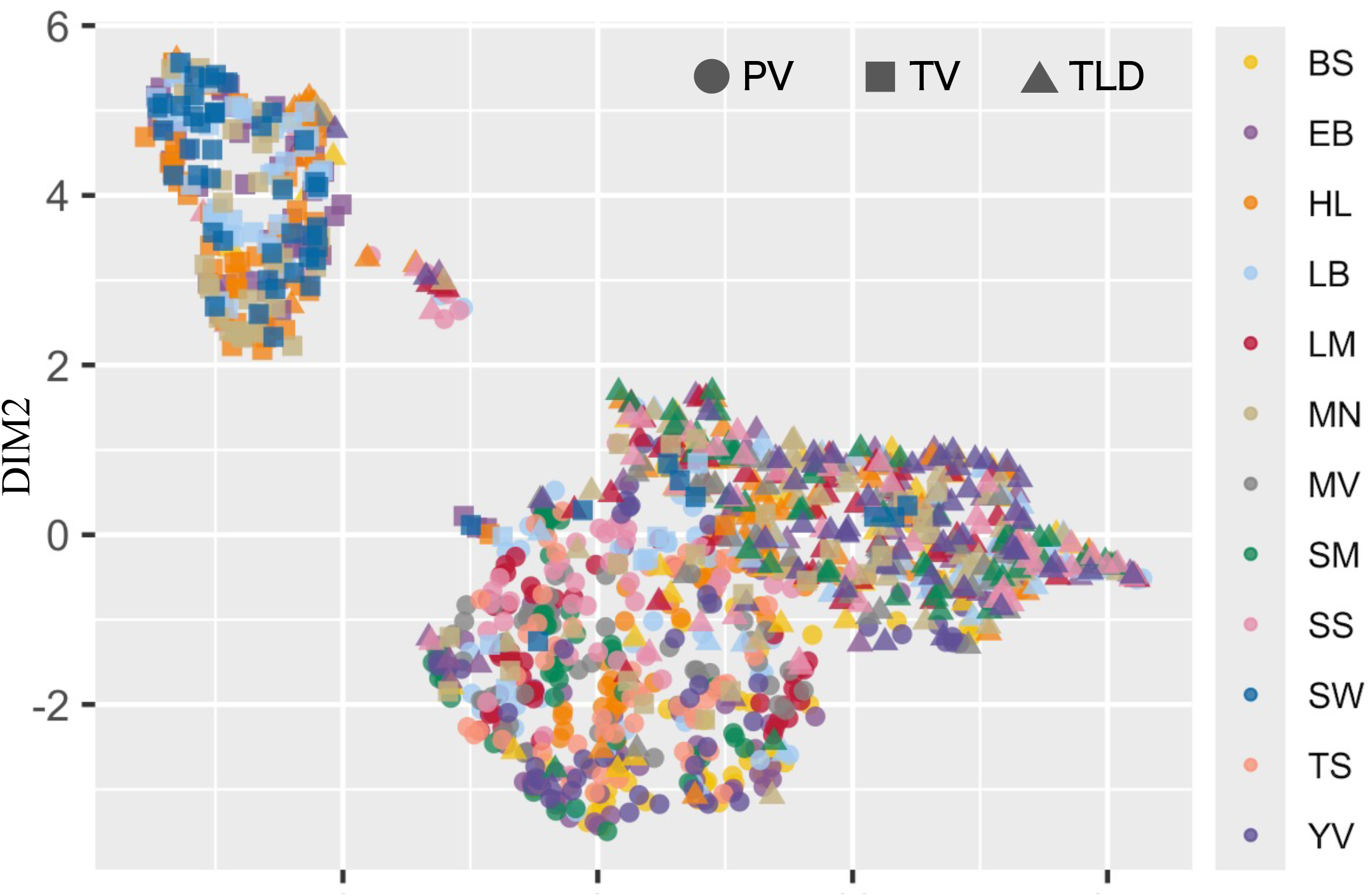
UMAP embedding result. DIM1 and DIM2 correspond to the first and second dimensions calculated by UMAP. Shapes represent different datasets, and colors denote disease types.

### Cross-dataset performance evaluation

Performance metrics, including precision, recall, and F1 score, are presented in Figures 2A–C. As expected, all models achieved high performance when training and validation were conducted on the same dataset. However, a pronounced decline was observed when models were validated on different datasets. For example, ResNet trained on PV, which consists of simple images with homogeneous backgrounds, achieved a high F1 score of 0.987 on PV validation. In contrast, validation on TV and TLD resulted in sharp drops to F1 scores of 0.012 and 0.130, respectively, despite TV also featuring isolated leaves captured under controlled conditions. Similarly, models trained on TV or TLD underperformed when validated on PV, illustrating the bidirectional nature of cross-dataset variability.

**Figure 2.**
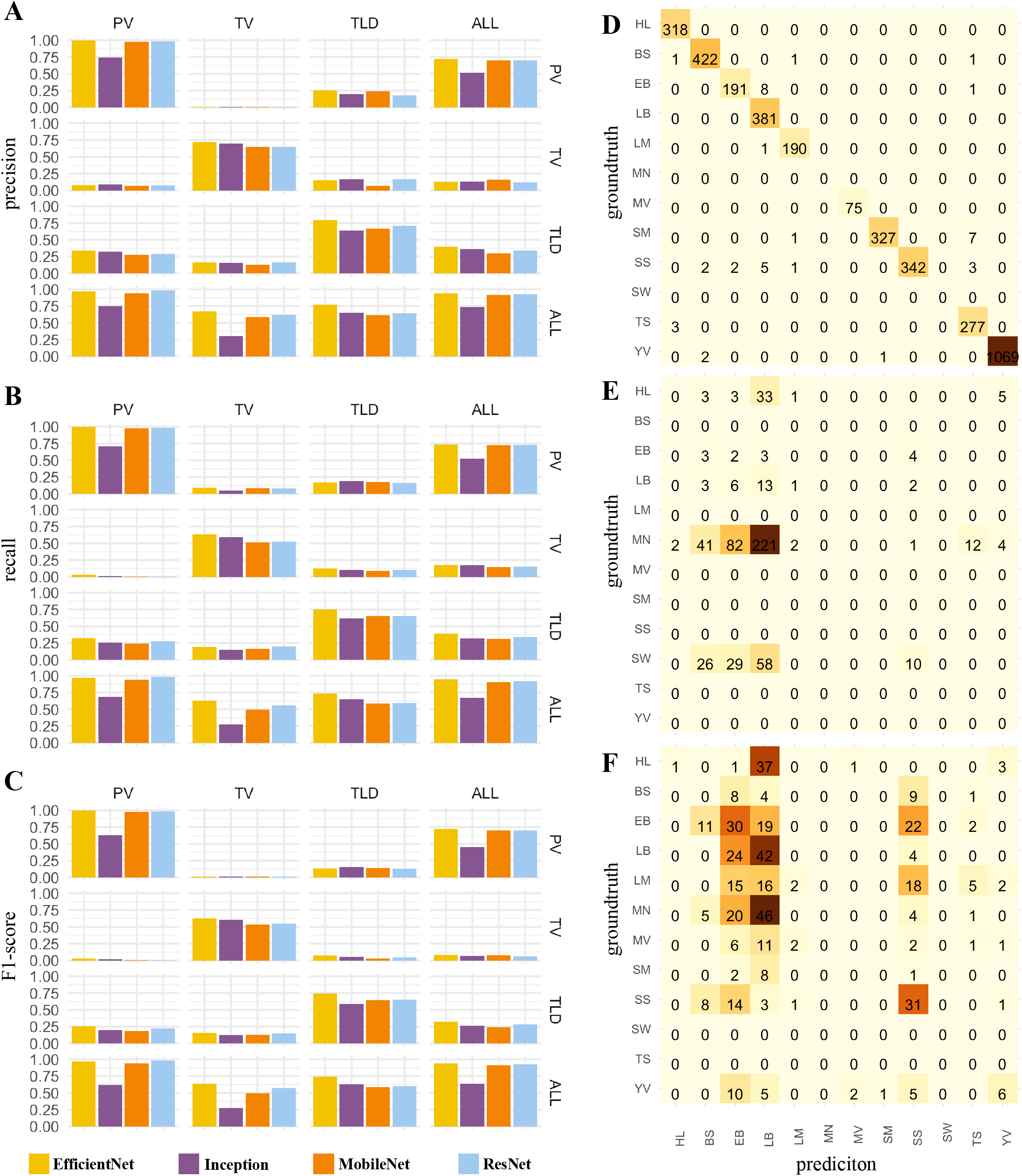
Performance metrics of plant disease classification models. Panels **A, B**, and **C** show the precision, recall, and F1 scores, respectively, for models trained on the datasets indicated in the rows and validated on those listed in the columns. Panels **D, E**, and **F** present confusion matrix heatmaps for ResNet trained on the PV dataset and validated on the PV, TV, and TLD datasets, respectively.

Training on a combined dataset (PV, TV, and TLD; collectively referred to as ALL) mitigated performance degradation to some extent, although performance still declined compared to intra-dataset validation. Performance was better preserved when validating on PV, likely due to the predominance of PV images in the combined dataset.

Confusion matrix analysis provided further insights. ResNet trained on PV reliably classified diseases in PV images (Figure 2D). However, when validated on TV, a large proportion (85.5%) of MN and SW images, absent from the training data, were misclassified, primarily as BS, EB, or LB (Figure 2E). Excluding MN and SW, performance remained poor (F1 score = 0.092), with frequent misclassification of HL images into BS, EB, LB, LM, and YV categories. Similarly, validation on TLD revealed comparable misclassification patterns, with MN frequently misclassified as EB or LB (Figure 2F). Even among categories common to TV and TLD, substantial misclassifications were observed. For example, HL images were often predicted as LB, and LM as EB, LB, or SS. Frequent misclassifications also occurred among EB, LB, and SS categories. A notable and consistent trend was the misclassification of HL images as LB (Figures 2E–F). This may result from physiological wilting in healthy leaves mimicking LB symptoms. Misclassification into other categories such as EB, LM, MV, SS, and TS was less frequent, likely because those diseases present more generalized symptoms across the entire leaf, unlike the localized wilting characteristic of LB.

Models trained on PV struggled with the complexity of field-collected images in TLD, which is expected given PV’s controlled imaging conditions. However, the poor generalization of models trained on TLD to PV images suggests that training with complex data alone is insufficient to address cross-dataset variability. These patterns were consistent across all tested architectures indicating that generalization challenges persist regardless of model architecture.

### Challenges in dataset availability and accessibility

Addressing cross-dataset variability requires access to larger, more diverse datasets. However, efforts to obtain additional tomato PPD datasets were hindered by limited availability. A survey of publicly available datasets revealed that many studies continue to rely on PV dataset for algorithm development. Field-collected datasets, which better reflect real-world agricultural variability, remain scarce and often inaccessible. Notably, despite frequent assertions in Data Availability statements that “datasets are available from the corresponding author upon reasonable request,” many requests went unanswered, even for recently published studies.

The creation and open dissemination of large, representative datasets would greatly advance the development of agricultural artificial intelligence (AI). Nonetheless, data privacy concerns and the high costs associated with data collection represent significant barriers to widespread data sharing. Even when datasets are compiled at considerable expense, making them freely available remains a challenge. Overcoming these obstacles and promoting the release of high-quality, field-representative datasets will be crucial for improving model generalizability and driving future innovations in agricultural AI.

### Concluding remarks

This study highlights the critical impact of cross-dataset variability on the performance of deep learning models for PPD diagnosis. Although models demonstrated high accuracy when trained and tested on the same dataset, their performance declined markedly when evaluated on different datasets, a trend consistent across all tested architectures. Additional analysis using flower species recognition tasks further confirmed this observation (SI Data). As generalization challenges cannot be mitigated merely by altering model architectures, future research should focus on the development and dissemination of diverse datasets. Addressing these challenges is essential for advancing the applicability of AI in agricultural research and practice.

### Limitations

A robust assessment of cross-dataset generalization in real-world settings necessitates large and diverse datasets that include images captured under actual field conditions. However, due to limitations in data availability and accessibility, this study was confined to four datasets, none of which were collected directly from agricultural fields. Future studies should incorporate more comprehensive evaluations as field-derived datasets become available.

## Declarations

### Availability of data and materials

The PlantVillage dataset is available at https://github.com/spMohanty/PlantVillage-Dataset. The Tomato-Village dataset is accessible at https://github.com/mamta-joshi-gehlot/Tomato-Village. The Tomato Leaf Disease dataset can be found at https://universe.roboflow.com/bryan-b56jm/tomato-leaf-disease-ssoha/dataset/63.

## Acknowledgments

Computational resources were partially provided by the SHIHO supercomputer at NARO.

